# Comparing the aftereffects of motion and causality across visual co-ordinates

**DOI:** 10.1101/499764

**Authors:** Regan M. Gallagher, Derek H. Arnold

**Affiliations:** School of Psychology, The University of Queensland, Brisbane, Australia.

**Keywords:** motion adaptation, causality aftereffect, perception, inference

## Abstract

Albert Michotte (1946/1963) introduced causality into the realm of experimental phenomenology. He disputed Hume’s (1739/1978) claim that the impression of causality comes only from conscious inference. Since then, causality adaptation studies have suggested that the visual perception of causality undergoes sensory adaptation, akin to the retinally-specific aftereffects of motion or orientation adaptation. Here we present 5 Experiments that, together, dispute a view of retinotopically-mapped neural populations dedicated to causality detection. We first point to key issues in previous studies of causality adaptation. Then we extend the basic causality adaptation paradigm to show that causality aftereffects occur in spatially global visual space. We directly compare causality aftereffects to the motion aftereffect to show important differences in their co-ordinate mapping. Our data point to a role for cognitive inferences as being an important aspect of causality aftereffects, despite causal impressions being tightly constrained by sensory perception.

---

Centuries ago, David Hume (1739/1978) argued that cause and effect must be inferred through a process of learned associations and cognitive inferences. His argument was that the relationship between two objects, or successive events, can only ever be correlated in the senses; sensory information is alone insufficient to determine causation in any objective way. Hume suggested that the impression of causality was inferred from experience, but never directly perceived.

Although Kant’s (1781) critique of pure reason tempered this strong empirical view, Hume’s general conclusion dominated Psychologists’ thinking on the topic until Michotte (1946/1963) introduced causality into the realm of experimental phenomenology. He acknowledged the essential truth of Hume’s claim that the mechanical information given to the senses provides no objective truth about cause. But Michotte, like Kant, disagreed with Hume’s presumption that the world, and more importantly the contents of the mind, consists of discrete objects and events. He pointed to the notion that perception and sensation are not direct replica of their physical stimuli. In a series of cleverly crafted experiments, he demonstrated the remarkable consistency and automaticity of subjective reports of causal impressions, which strongly depended on sensory factors. On this basis, Michotte argued that the causal impression (see Figure 1) is a direct phenomenal experience in the same way we perceive motion and colour and form.

**Figure 1.**
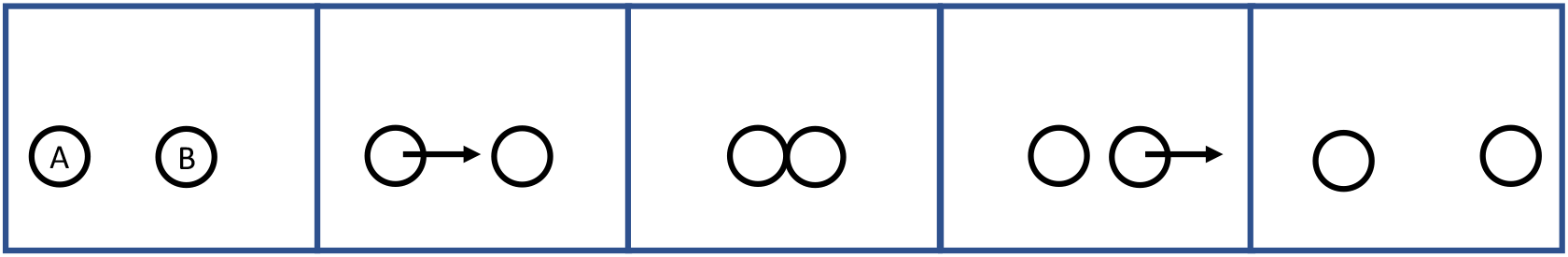
A standard launch event. The first object, A, moves from the left towards the stationary second object, B. Upon contact, object A becomes stationary and object B begins moving along the same trajectory. Participants typically report that A was the cause of B’s movement.

Critics of Michotte’s conclusion point out that his studies relied heavily on subjective reports (see Wagemans, van Lier, & Scholl, 2006; Choi & Scholl, 2009). Typically, Michotte’s participants would view the apparent interaction of objects, and then verbalise or record their experience by writing down their subjective impression. This methodology leaves open the influence of cognitive inferences in describing experience. Nevertheless, Michotte’s work had a profound impact on the view of causality as a perceptual phenomenon. Part of the remaining challenge for distinguishing causal impressions as perceptual or cognitive is to use quantitative methods over subjective reports.

## Does causality adapt?

Sensory adaptation and psychophysics allow for a principled empirical approach to studying the neural computations underlying perception. Aftereffects have become invaluable tools in perception research — this is summarised by the Psychologists’ refrains that “adaptation is the psychologist’s micro-electrode” and “if it adapts, it exists”. Measuring the parameters of perception using aftereffects allows us to estimate, often with high precision (Van Wezel & Britten, 2002), the degree to which sensory adaptation impacts stimulus-selective neural populations. In this way, recent work (Rolfs, Dembacher, & Cavanagh, 2013) provides an important advancement to Michotte’s legacy. In their experiments, Rolfs and colleagues applied the tools of sensory adaptation to test for a perceptual basis of causal impressions.

Rolfs and colleagues’ (2013) experimental manipulation is an elegant and intuitive test of perceived causality. In their study, participants repeatedly viewed a contact-launch scenario in which object A moved toward object B and, upon contact (but without overlap), object A stopped moving and object B moved at the same speed along the same trajectory. Then, participants were presented with a test scenario: the degree of spatial unity between A and B was systematically varied so that sometimes the two made only contact, and other times A overlapped B. Participants categorised each event as either a non-causal pass (if object A appeared to pass through object B) or a causal launch (if object B’s motion appeared causally initiated by object A).

Results of the experiments suggested that causality responses became less likely following repeated collision events; participants were more likely to report previously ambiguous events as non-causal passes. One of the critical manipulations was to test for causal impressions in the same location as the adaptation events versus an unadapted location. The experimenters found that the reduced propensity to see test events as causal was (largely) limited to events in retinotopic locations. Such a conclusion agrees with and extends the conclusions of Michotte, that causality is a perceptual phenomenon. There are, however, several reasons to be sceptical of the interpretation that adaptation of the visual perception of causality is retinotopically mapped.

## Is the reference frame of causality aftereffects retinotopic?

An important finding of Michotte’s work, although perhaps not so well-known, is that the impression of causality could be induced even in the absence of a physical connection between the first and second object (termed ‘action at a distance’; Michotte, 1963; p.p. 99-100). His experiments showed that spatial unity was not a necessary condition for producing causal impressions using a launch paradigm. Rather, Michotte found that causal impressions were recovered from stimuli with spatial separation, and launch impressions decreased when the disunity between objects increased. This finding has important implications for how best to measure and interpret the impression of causality (see Figure 2).

**Figure 2.**
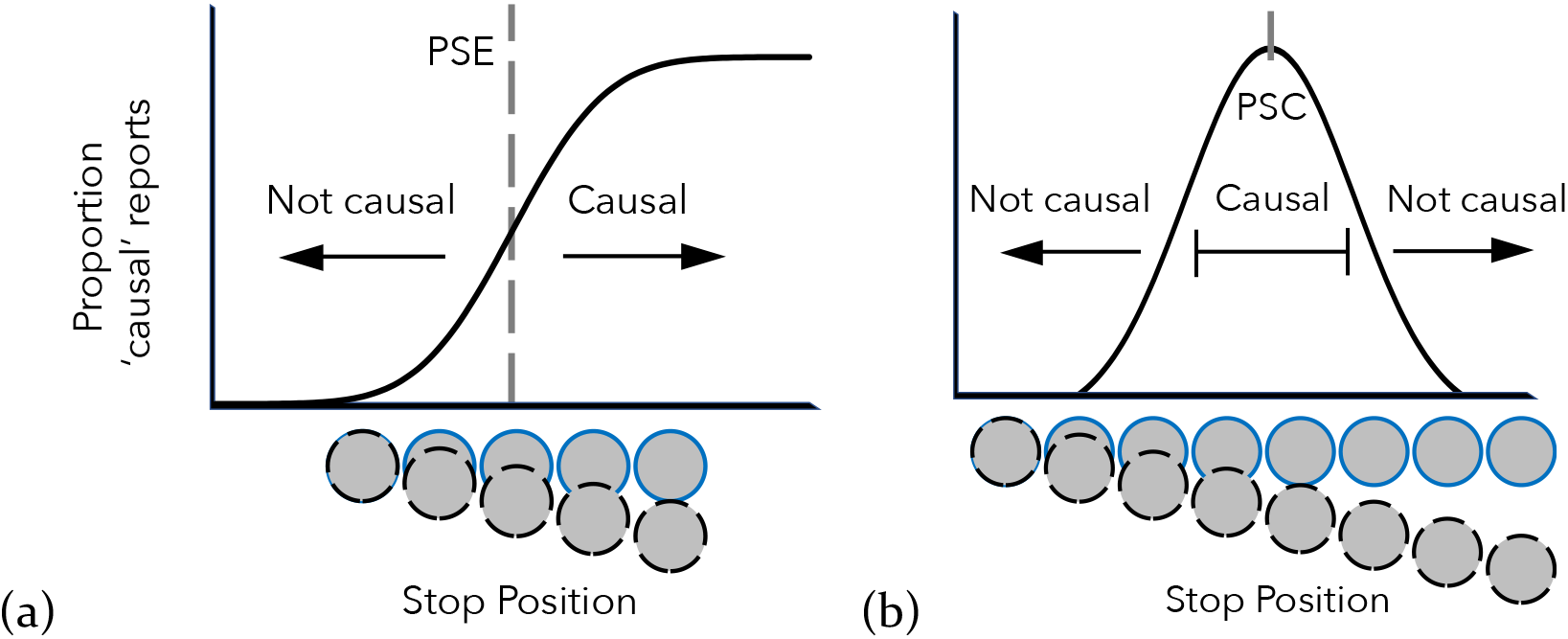
What can we infer from causality judgements? Fitting a cumulative Gaussian function across spatially coincident stimuli (panel a) estimates the point of subjective equality (PSE) between ‘causal’ and ‘non-causal’ impressions. Fitting the raised Gaussian curve (panel b) provides an estimate of the point of subjective causality (PSC). This model suggests that causal impressions are experienced between two spatial boundaries, within which the interaction of two stimuli can lead to reports of a causal impression. The raised Gaussian curve also predicts that a broader range of stimuli can be subjectively causal, with some causal impressions arising in the absence of spatial connection.

In causality adaptation studies, the perceptual qualities of interest have often [1,2,3] been modelled using a cumulative Gaussian function (see Figure 2a), rather than a Gaussian function (see Figure 2b). These models measure similar processes; for example, the parameters of each model provide estimates for the physiological sensitivity and accuracy of the information encoded by the observer’s sensory system(s). The two models also have similar underlying assumptions of normality, but when applied to causality judgements, might provide different estimates of central tendency and precision.

Rolfs and colleagues (2013) argued that causality adaptation resulted in a fatigue-like neural process, with all the hallmarks of sensory adaptation. They suggested that low-level, retinotopically organised causality detectors became less likely to encode causality after being adapted to causal interactions. While changes in cognitive criteria could account for a generally higher response threshold after repeatedly seeing instances of causation (Powesland, 1959), it is unlikely that a cognitive explanation can account for strict retinotopic mapping. However, if causality judgements are better described by a standard normal distribution, rather than the cumulative normal distribution used in their experimental modelling, then Rolfs and colleagues (2013) selected their adapting stimulus from a value at or near the central tendency of the subjective causality distribution — in effect adapting to a neutral’ stimulus. Indeed, extending Rolfs and colleagues data to describe a larger distribution of causality judgements suggests this prediction is plausible (see Figure 3). This could have consequences for the interpretation of changes in causality measurements.

**Figure 3.**
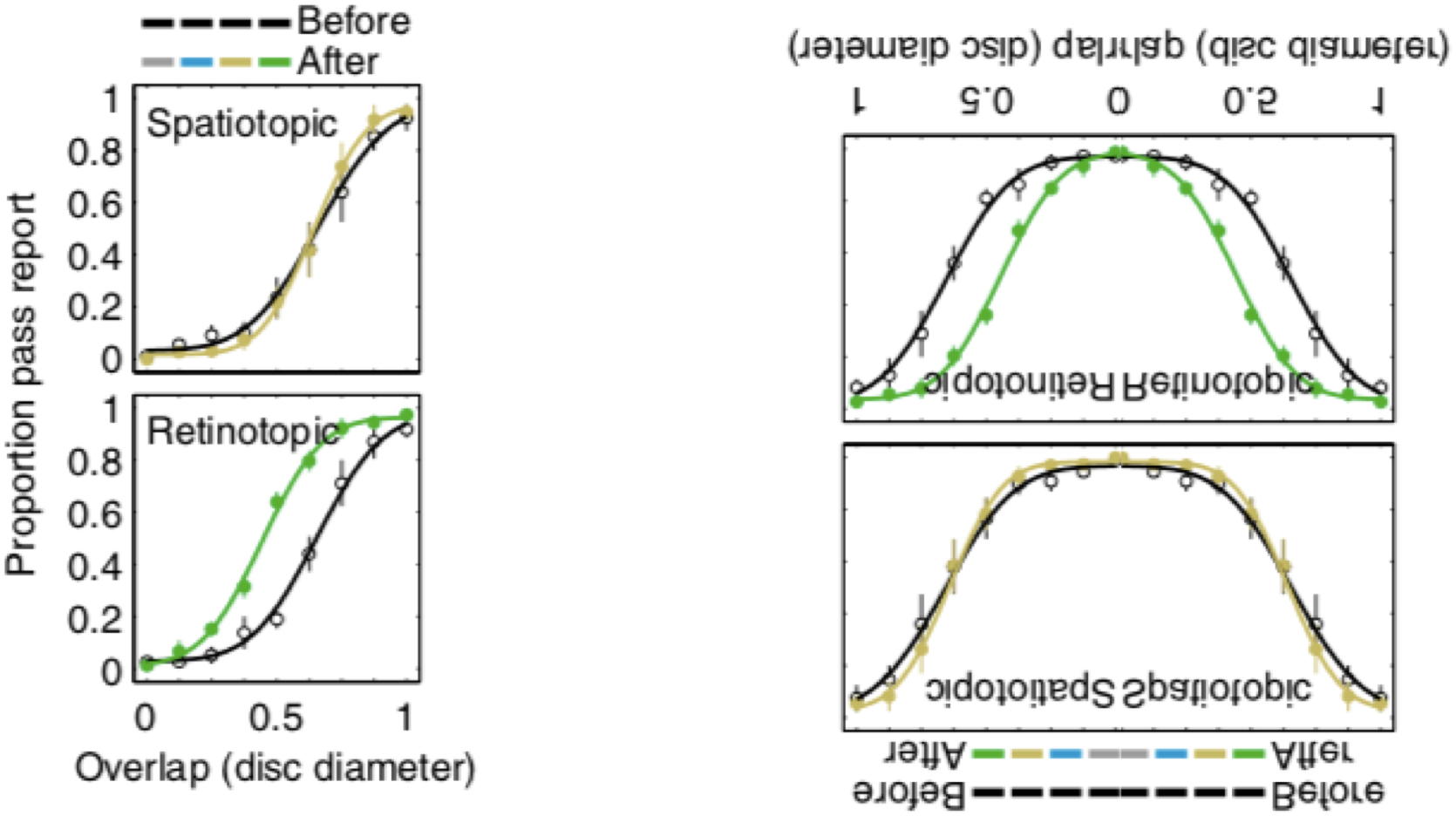
Results of adaptation in Rolfs et al.’s experiments (left), rotated and mirror flipped (right). These data create a prediction for what should happen in each condition when measured across the broader causality distribution. These data predict no shift in central tendency when the adapting stimulus is a neutral stimulus.

**Figure 4.**
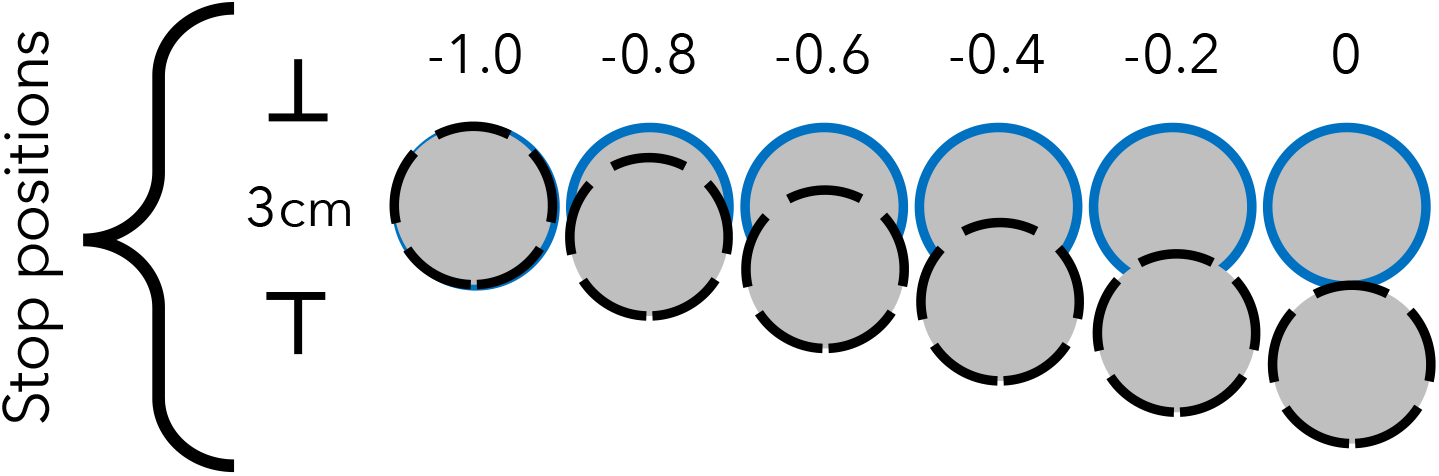
Contact-launch stimuli where each number refers to the stopping location of the first disc. Depicted here is the range of test overlaps used in Experiment 1. A stopping position of o refers to no disc overlap (full contact) and −1.0 means the first disc stopped when it had fully overlapped the second disc (complete pass).

## The present study

In the present study, we argue against a retinotopically mapped feature detection view of causality. We first suggest that Rolfs et al.’s (2013) data could equally be explained by a change in expectation of future interactions based on past experience. Consistent with this perspective, we present data that suggests causality aftereffects (measured by central tendency changes) are a spatially global phenomenon, rather than retinotopically mapped. We adopt the overall paradigm pioneered by Rolfs and others, as we recognise it has great benefit for understanding the neural computations underlying causality judgements. The present study addresses potential limitations of a restricted test range. We conceptually replicate the causality adaptation paradigm and use an extended range of launch stimuli to include a range of non-contiguous stimuli as used in Michotte’s early experiments.

We hypothesise from previous adaptation studies that, if causality is retinotopically mapped, changes in the central tendency of causal impressions should change only in the adapted region. If fitting a model to the restricted test range has no impact, this should be equally true for the full distribution. However, if adaptation aftereffects arise from expectations, we might not expect to see a strict retinotopic mapping of causality adaptation (see Arnold, Petrie, Gallagher, & Yarrow, 2015).

To test these questions, we first replicated the causal launch effect across the range used in previous studies of causality adaptation. To this data we fit two inferential models (cumulative and raised Gaussian functions) to compare estimates of central tendency and precision. Then, we extend previous studies by measuring subjective causality across a broader range of causal launch scenarios. With an extended stimulus range we show that, while causality adaptation produces reliable decision aftereffects, our experiments directly repudiate an interpretation of retinotopically mapped causality detectors. We therefore find further support that causality aftereffects result from an expectation about how objects should behave in a collision (Arnold et al., 2015), rather than a change in a strictly retinotopic mapping of the perception of causality (Rolfs et al., 2013), and thereby distinguish causality aftereffects from classic sensory adaptation.

## Experiment 1

Previous studies leave open the possibility that causality adaptation was modelled using a restricted stimulus range and interpreted using unsuited function parameters. This ambiguity has important implications for how the impact of causality adaptation should be interpreted. In Experiment 1 we replicate the range of stimuli measured in previous studies and then fit two separate models, the cumulative and raised Gaussian functions. We compare the estimated parameters of subjective causality based on these models, and show that these models lead to different conclusions.

### Method

#### Participants

Ten participants from the University of Queensland volunteered their time in exchange for course credit as part of a first-year Psychology participation scheme. All were naïve to the purpose of the experiment. Each participant provided verbal consent, acknowledging they could cease participation at any time and for any reason without prejudice or penalty. Ethical approval for all experiments was obtained from the University of Queensland’s Ethics Committee.

#### Materials & Stimuli

Stimuli were presented on LCD monitors running Windows XP. All computers were running Matlab software and the Psychophysics Toolbox (Brainard, 1997; Pelli, 1997). All monitors had a screen refresh rate of 60Hz. Test stimuli were two identical circles each approximately 3cm in diameter. Stimuli moved across the screen at a rate of 0.10 diameters per frame (6 diameters per second) for a minimum of 12 diameters and a maximum of 15 diameters. The starting location of the first disc and the stopping location of the second disc were each varied by up to 1.5 disc diameters at random on a trial-by-trial basis. Disc A was initially stationary and always began moving first, Disc B was stationary until Disc A stopped moving. Disc A travelled in a linear path until the stopping location, then upon stopping, the motion transferred from Disc A to B, and Disc B continued along the same linear trajectory.

The temporal relationship between Disc A stopping and Disc B moving was held constant, so that Disc B always launched into motion on the frame immediately after Disc A stopped. The spatial relationship between the two discs at the time of launch was systematically varied. At maximum, Disc A entirely occluded Disc B before stopping (stop position of −1.0). At minimum, Disc A stopped moving when it contacted Disc B, but had no spatial overlap (stop position of 0.0). The remaining overlaps were linear steps of 0.20-disc diameters (see Figure 3).

#### Instructions

Participants were instructed to report whether Disc A appeared to make contact with *and* launch Disc B into motion by pressing the right arrow (causal response) on the keyboard. If it appeared as though Disc A moved entirely through Disc B without making contact, or if Disc B moved independently of Disc A’s motion, participants were instructed to press the left keyboard button (non-causal response).

#### Procedure

Participants sat comfortably in a chair approximately 55cm away from the display, rested their hands on the keyboard’s directional buttons, and fixated a central cross-hair. Disc A and Disc B appeared on screen, and Disc A began moving after a random inter stimulus interval (ISI) of not more than 1500ms. Disc A moved toward its stopping location and, upon arrival, the motion of Disc A was transferred to Disc B, which continued moving until it reached its stopping location. When Disc B stopped, participants were prompted for a response by a flickering of the fixation cross from white to black and back to white. When the participant responded, a new trial began immediately following the ISI. Each of the six stimulus values were repeated 50 times, so that all participants attended 300 trials in total. The experiment lasted approximately 25 minutes.

#### Data analysis

Data were analysed using GraphPad PRISM software (Prism 7, 2017). The Nonlinear regression toolbox was used to fit all psychometric functions. Individual fits were calculated using automatic outlier elimination, and outlying or missing data were interpolated using a 95% confidence interval. For each dataset, we compared which of the two models (cumulative or raised Gaussian) provided the best fit to the data. All parameters were unconstrained (2 free parameters for the cumulative Gaussian, 3 free parameters for the raised Gaussian). We obtained likelihood estimates using Akaike’s Informative Criteria (AIC) comparison, and generated null hypothesis significance test values using an Extra Sum-of-Squares *F*-test.

### Results

We first checked that subjective causality can be equally modelled using both the cumulative and raised Gaussian functions. Cumulative Gaussian functions were fit to each participant’s distribution of causality judgements, and 50% points were taken as estimates of the point of subjective equality (PSE) — the stimulus value equally likely to be judged as causal and non-causal. A standard normal Gaussian function was also fit to each individual’s distribution of causality responses, and the central tendency of the fitted function was taken as an estimate of the point of subjective causality (PSC).

#### Model comparisons

To demonstrate the effect of fitting different models to the same causality data, we estimated the PSE and PSC from participants’ responses using both cumulative and raised Gaussian functions, respectively. The Extra Sum-of-Squares F-test shows that the parameters of the two distributions provided marginally different estimates of central tendency (cumulative Gaussian fit: *M* = −0.43, *SD* = 0.62; raised Gaussian fit: *M* = −0.15, *SD* = 0.46; difference: *F*_1,57_ = 3.97, *p* = .051). Assessing the goodness of fit of each model using the Akaike Informative Criteria (AIC) comparison, suggests that the raised Gaussian function provides an overall better fit than the Sigmoid function (Table 1).

**Table 1.** Comparison of model fits.

| Model | R<sup>2</sup> | df | absSS | Likelihood |
| --- | --- | --- | --- | --- |
| Gaussian function (PSC) | .46 | 57 | 3.21 | 70.43% |
| Sigmoidal function (PSE) | .42 | 58 | 3.44 | 29.57% |

*Note.* Goodness-of-fit model parameters reported.

In previous studies, the adapting stimulus had an overlap value of 0%. If the causality distribution is best described by a raised Gaussian function, this value might be a ‘neutral’ stimulus (at the peak of the function). We therefore tested whether the central tendency of each function differed from 0% overlap. An Extra Sum-of-Squares *F*-test showed that the central tendency of the cumulative Gaussian function was significantly different to 0% disc overlap (*F*_1,58_ = 20.49, *p* < .001). The central tendency of the raised Gaussian function, however, was not significantly different to 0% overlap (*F*_1,57_ = 1.15, *p* = .288).

#### Decision auto-correlation

To demonstrate that changes in the PSE and PSC can lead to different conclusions about the underlying distribution of subjective causality, we conducted a decision auto-correlation test on causality judgements and measured the effect of prior decisions on these parameter estimates. The correlation between each individual’s response on consecutive trials provides an indication of how much the previous response impacts the next response (independent of the physical stimulus). Each participant’s data was binned according to their response on the last trial (causal or non-causal), and the two psychophysical models (cumulative and raised Gaussian) were again fit to participants’ data. Responses were then analysed for trials following a subjective report of ‘causal’ and compared against trials following a subjective report of ‘non-causal’ (see Figure 5).

**Figure 5.**
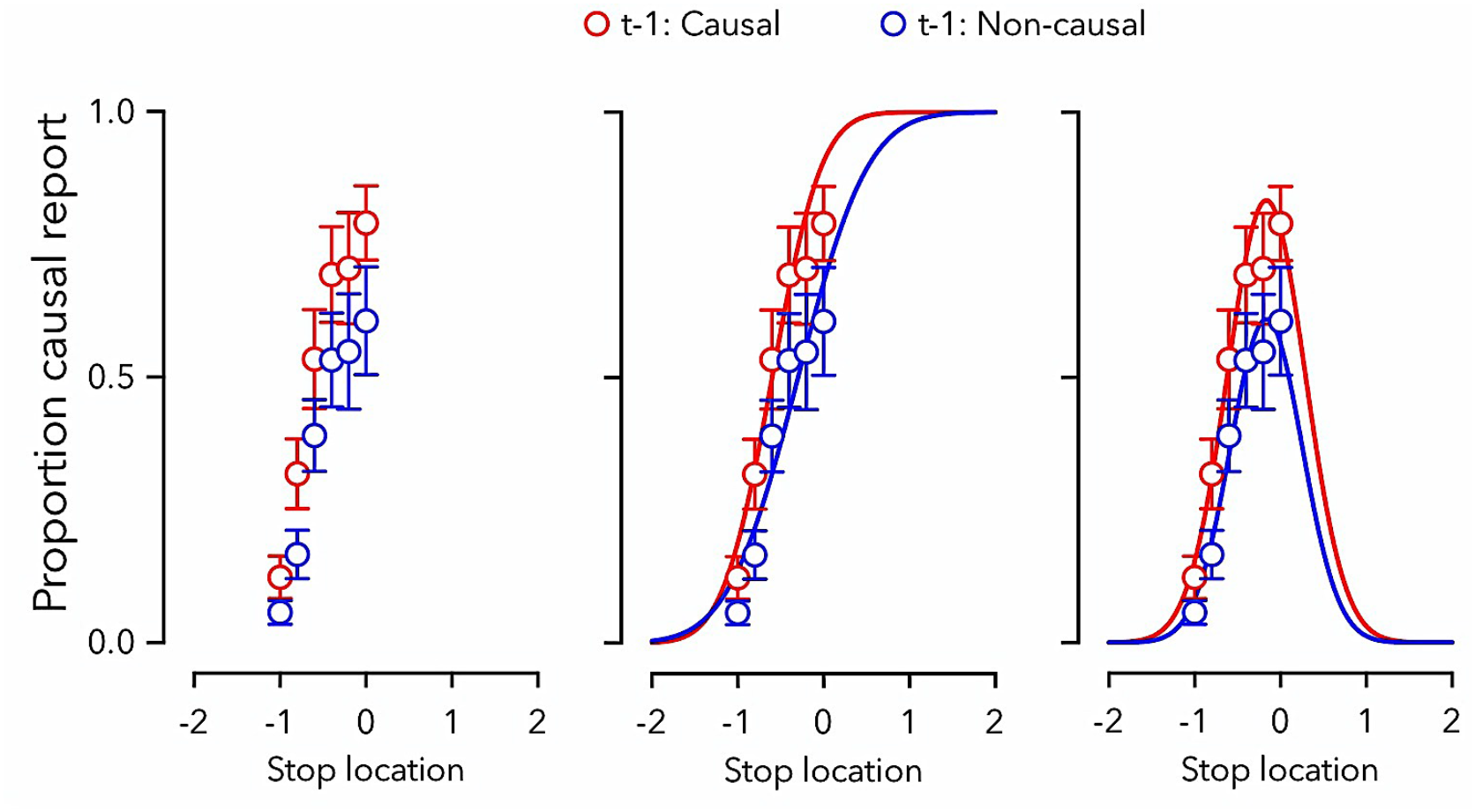
Contact-launch data where 0 refers to no disc overlap (full contact) and −1 means full disc overlap (complete pass). Depicted here is a test of decision auto-correlation, where data are binned according to the response on trial t-1 (causal or non-causal). The same data shown on the left in (a) are fit with (b) a cumulative Gaussian function or (c) a raised Gaussian function. The different models lead to different interpretations for the effect of prior decisions on the central tendency of function fits, and each model fit makes different predictions about the patterns of data points presently unmeasured across the stimulus continuum.

**Figure 6.**
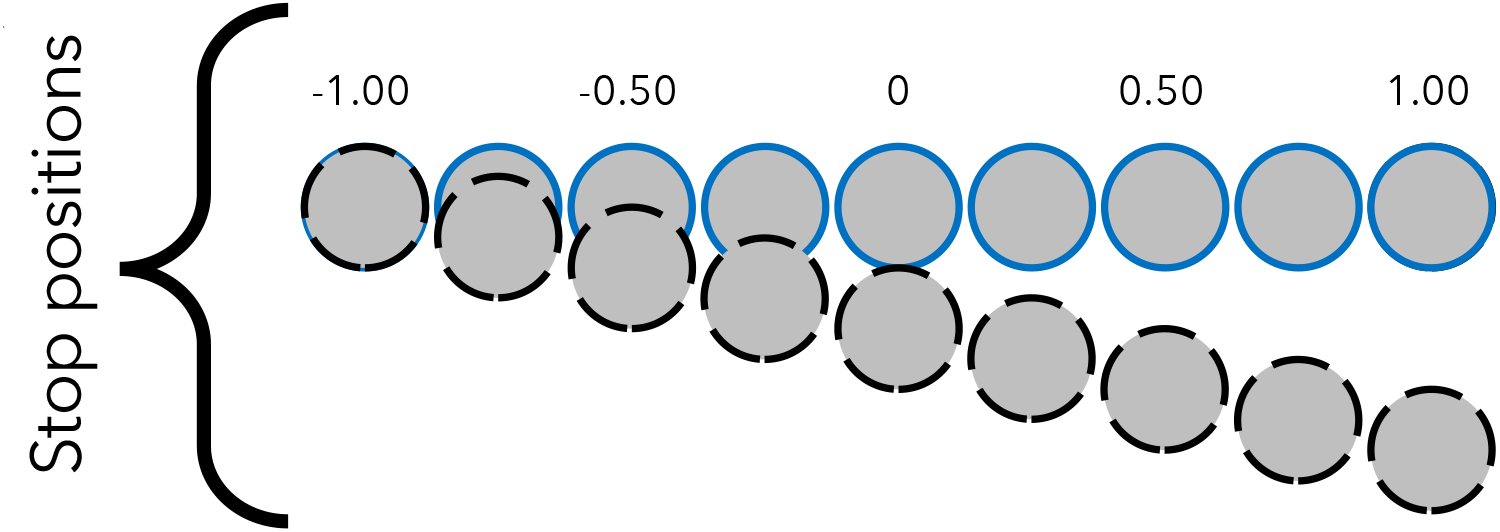
Launch stimuli used as test events in Experiments 2–4. The adapting event had a stop position of 0.0.

Fitting the cumulative Gaussian model suggested that the previous decision impacted the central tendency (causal: *M* = −0.60; non-causal *M* = −0.30; *F*_1,114_ = 17.02, *p* < .001) but not the standard deviation (causal: *SD* = 0.45; non-causal *SD* = 0.64; *F*_1,114_ = 1.90, *p* = .171) of the psychometric function. Fitting a raised Gaussian, on the other hand, had no effect on the central tendency (causal: *M* = −0.18; non-causal *M* = −0.14; *F*_1,112_ = 0.12, *p* = .735) of the psychometric function, but instead impacted the dispersion around the mean (causal: *SD* = −0.44, Amp = 0.84; non-causal *SD* = −0.43, Amp = 0.61; *F*_2,112_ = 4.26, *p* = .017).

### Discussion

Results of Experiment 1 indicate that the chosen model for describing the distribution of causal impressions can lead to different conclusions about the psychological processes that determine subjective causality. A simple decision bias — in this case, participants were on average more likely to repeat a previous response — can translate the central tendency of a cumulative distribution without impacting the central tendency of a raised Gaussian function, even when they describe the same data. To test whether fitting the incorrect model could have impacted previously reported causality adaptation aftereffects, the following Experiments examine a broader range of causality stimuli and the effect of launch adaptation on causal impressions.

## Experiment 2

Fitting the different psychophysical models indicated that the raised Gaussian function could be the preferred model for causality judgements. However, if this is the preferred model, only one-half of the relevant distribution was sampled in Experiment 1 (and previous studies). Extrapolating from the cumulative and raised Gaussian functions provide clear predictions for how launch events should be interpreted across a broader distribution of spatial contingencies. Both models predict that causal impressions should be recovered despite a lack of physical coincidence. Only the raised Gaussian model suggests that causal impressions should decay with both increasing spatial disunity and with increasing spatial overlap. To empirically determine which function best describes causality judgements, we replicate the contact-launch adaptation protocol pioneered in Rolfs and colleagues’ (2013) study using a broader range of launch events.

### Method

Experiment 2 was identical to Experiment 1, except for the following aspects. Participants completed two testing blocks: one baseline block, which was identical to Experiment 1, and one adaptation block, which involved passive viewing of repeated contact-launch events prior to each causality judgement. On the first trial in the adaptation block, participants witnessed 18 collisions prior to the test event, and on each subsequent trial participants witnessed 6 collisions prior to each test event. In total, 15 participants completed both the baseline and adaptation phase of Experiment 2. Four of the nine launching stimuli involved some degree of spatial separation. In both the baseline and the adaptation conditions, participants viewed 15 launch events for each of 9 spatial contingencies (stop positions). These events ranged from a stop position of-1.0 (complete overlap) through 0.0 (contact-launch) to 1.0 (one diameter separation) in steps of 0.25 diameters (see Figure 5).

### Results

We first examined whether or not extending the range of test stimuli (to include spatial disunity at the moment of launch) results in a decay of causal impressions. We fit a raised Gaussian function with 3 free parameters, describing the mean, standard deviation, and amplitude of the response distribution across the range of tests.

#### Baseline

Baseline data were well-described by the raised Gaussian distribution (Goodness-of-fit: R^2^ = .70). We next checked whether launch responses were likely to occur from stimuli with spatial disunity at the moment of launch. The stimulus most likely to appear causal at baseline had 22% overlap (95% Cl: 18% - 27%) between Disc A and B at launch. The standard deviation of the response distribution was 38% of a disc diameter (95% CI: 34% - 42%). The width of this function demonstrates that stimuli with spatial separation at launch can result in causal impression, and this impression decreases as a function of spatial disunity (see Figure 7, top left panel).

**Figure 7.**
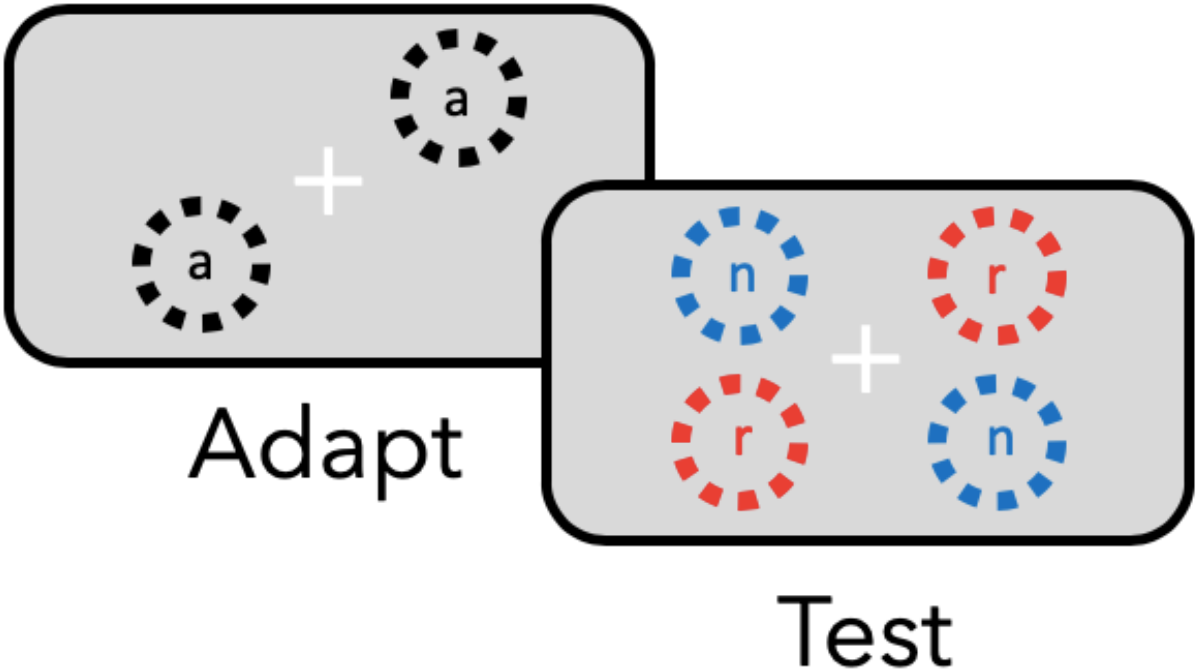
The co-ordinate mapping between adapting stimuli (a) and retinotopic (r) and non-specific (n) tests in Experiments 3–5. Adapting and test events were presented in two locations simultaneously to mitigate reflexive eye movements away from fixation.

**Figure 8.**
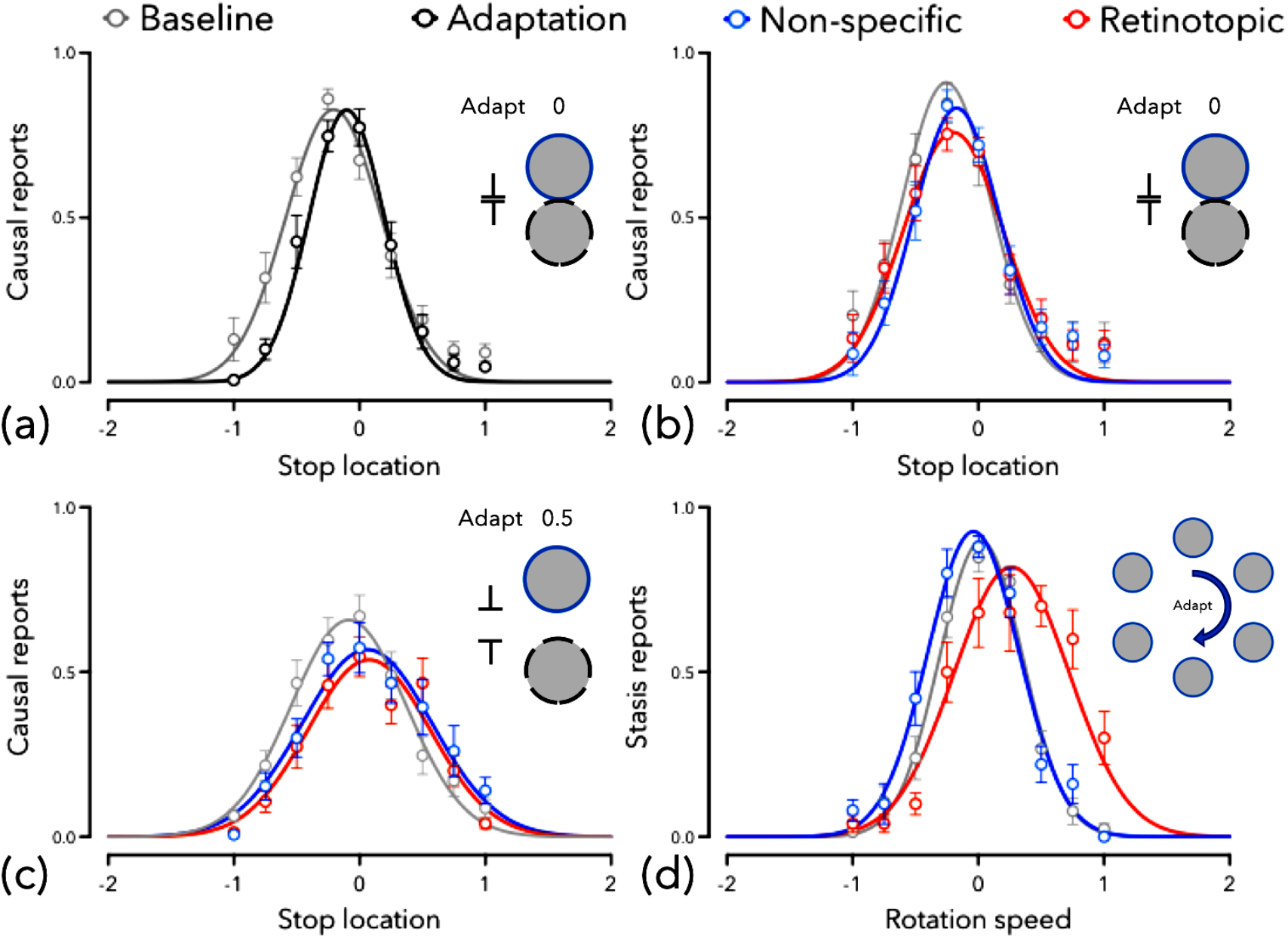
Results of adaptation in Experiments 2–5. (a) Experiment 2, (b) Experiment 3, (c) Experiment 4, (d) Experiment 5.

#### Effect of adaptation

We then examined the effect of adaptation to the contact-launch stimulus. We found that adaptation to 0% spatial overlap strongly impacted the distribution of causality responses *F*_3,348_ = 7.80, *p* < .001). Further evaluation showed that the central tendency of causality responses shifted towards the adapting stimulus, so that the most likely stimulus to evoke a causal response had 10% disc overlap (95% CI: 7% - 13%; *F*_1,258_ = 18.02, *p* < .001), and the standard deviation of the distribution was reduced to 30% of a disc’s diameter (95% CI: 27% - 33%; *F*_1,128_ = 9.29, *p* = .003).

### Discussion

Results of Experiment 2 show that causality judgements are best described by a raised Gaussian distribution. Causal impressions were reported for stimuli with spatial disunity at launch, but this impression rapidly decayed with increasing spatial separation. Surprisingly, we also found that adaptation to 0% disc overlap produced a shift in the central tendency of the causality distribution. However, this could be the result of increased precision following adaptation (i.e., repeated examples of causal launching create a more precise expectation). Experiment 3 tests the co-ordinate mapping of causality adaptation, which could indicate whether response changes likely arose from changes to perceptual or decision processes.

## Experiment 3

Adaptation of low-level causality detectors remains a plausible interpretation of the data in Experiment 2. It also remains plausible, however, that prolonged and repeated viewing of causal launches changes decision criteria; viewing repeated launches could have the effect of making an observer more discerning about what constitutes a causal interaction. Experiment 3 addresses this ambiguity by measuring the co-ordinate mapping of the adaptation aftereffect. To rule out changes to decision processes as an explanation, Experiment 3 compares the effect of adaptation across retinotopic and non-retinotopic test locations.

### Method

The Methods for Experiment 3 were the same as Experiment 2, except that tests appeared in either the same location as the adapting stimulus (retinotopic) or in a different location (non-specific). We also tested a new sample of 15 naïve first year Psychology students.

### Results

#### Effect of adaptation

We examined the effect of adaptation to the launch stimulus and found adaptation had a general impact on the distribution of causality responses (baseline: *M* = −0.24, *SD* = 0.34, Amp: 0.88; adaptation: *M* = −0.16, *SD* = 0.37, Amp 0.78; difference: *F*_3,223_ = 3.84, *p* = .010). Further evaluation showed that adaptation shifted the central tendency of causal impressions, so stimuli with spatial disunity were more likely to be judged causal, and stimuli with spatial overlap were less likely to be judged as causal after adaptation (*F*_1,220_ = 10.22, *p* = .002). We observed no significant changes in the standard deviation (*F*_1,224_ = 0.55, *p* = .461) following adaptation.

#### Co-ordinate mapping

We then examined whether the effect of adaptation was dependent on the location of the test stimulus. Results showed that the impact of adaptation changed the central tendency of retinotopic tests (*M* = 0.18, *SD* = 0.39, Amp: 0.73; baseline vs retinotopic: *F*_1,223_ = 3.96, *p* = .048) and also changed the central tendency of the non-specific tests (*M* = 0.14, *SD* = 0.34, Amp: 0.82; baseline vs non-specific: *F*_1,223_ = 11.95, *p* = .001). However, comparing test locations showed no statistical difference in central tendency (retinotopic vs non-specific: *F*_1,228_ = 0.99, *p* = .320). These results show that, after adaptation, causal impressions became more likely for events with spatial disunity, and less likely for events with spatial overlap, and this was true for both retinotopic and non-retinotopic tests.

### Discussion

Results of Experiment 3 suggest that causality aftereffects are location independent. Tests in the same location, as well as tests in a different location, both demonstrated a shift in central tendency after repeated presentations of causal launching. Moreover, these results could not be explained solely by a shift in the standard deviation of the response distribution. However, adaptation to a near-neutral stimulus could be mitigating the adaptation aftereffect. Experiment 4 addresses this possibility by testing for adaptation to launch events with spatial disunity (i.e., using a non-neutral adapting stimulus).

## Experiment 4

It remains an open possibility that the impact of launch adaptation is constrained by the use of a ‘neutral’ contact-launch stimulus. Such a constraint might limit the effect of causality adaptation, reducing power for detecting retinotopically-mapped adaptation of causality perception. Experiment 4 extends the adaptation paradigm to test for an aftereffect to launch events that have spatial disunity. We again tested for a location dependent effect of launch adaptation.

### Method

The Methods for Experiment 4 were the same as for Experiment 3, except that Disc A of the adapting stimulus stopped short of Disc B by 0.5 diameters. We also tested a new sample of 15 naïve first year Psychology students.

### Results

#### Effect of adaptation

We examined the effect of adaptation to spatial disunity in a launch stimulus. We found that adaptation strongly impacted the distribution of causality responses (baseline: *M* = −0.11, *SD* = 0.44, Amp: 0.68; adaptation: *M* = −0.04, *SD* = 0.47, Amp 0.57; difference: *F*_3,264_ = 5.47, *p* = .001). Further evaluation showed that adaptation shifted the central tendency of causal responses, so stimuli with spatial disunity were more likely to be judged causal, and stimuli with spatial overlap were less likely to be judged as causal after adaptation *F*_1,264_ = 11.55, *p* < .001). We observed no changes in the standard deviation *F*_1,264_ = 0.48, *p* = .488) following adaptation.

#### Co-ordinate mapping

We then examined whether the effect of adaptation was dependent on the location of the test stimulus. Results showed that the impact of adaptation had a clear impact on the central tendency of retinotopic tests (*M* = 0.07, *SD* = 0.48, Amp: 0.54; baseline vs retinotopic: *F*_1,264_ = 11.21, *p* = .001) and also a clear impact on the non-specific tests (*M* = 0.06, *SD* = 0.52, Amp: 0.57; baseline vs nonspecific: *F*_1,264_ = 8.67, *p* = .004). Comparing test locations showed no statistical difference in central tendency (retinotopic vs non-specific: *F*_1,264_ = 0.61, *p* = .436). These results show that, after adaptation, causal impressions became more likely for events with spatial disunity, and less likely for events with spatial overlap, and this was true for both retinotopic and non-retinotopic tests.

### Discussion

Results of Experiment 4 show that non-overlapping adapting stimuli can still produce a causality adaptation aftereffect, and this effect was demonstrated to have location independence. The impact of adaptation on both retinotopic and non-retinotopic locations indicates that inferences of causality are likely not driven by low-level, retinotopically-mapped causality detection mechanisms. Unless, of course, our Experiments could not dissociate retinotopic from non-retinotopic adaptation effects, even if they exist. Experiment 5 addresses this question by using motion adaptation rather than causality adaptation.

## Experiment 5

It remains to be determined that our Experimental design can detect a retinotopic adaptation aftereffect. For example, if participants made reflexive eye-movements towards each adapting and test stimulus, then each test would be retinotopically mapped regardless of spatial location. While this was controlled for by having symmetrically matched test presentations, we wanted to ensure that this did not impact our results. We therefore tested whether our paradigm could detect a retinotopic mapping using an effect with a known retinotopic reference frame (Knapen, Rolfs, & Cavanagh, 2009). To this end, Experiment 5 tests the co-ordinate mapping of the motion aftereffect using the same paradigm used here for causality adaptation. If participants were reflexively looking at both the adapting and test stimuli, we should observe equivalent motion aftereffects across retinotopic and non-specific test locations.

### Results

#### Effect of adaptation

We examined the effect of adaptation to a rotating motion stimulus and found that adaptation to clockwise rotation strongly impacted the distribution of stasis responses (baseline: *M* = 0.02, *SD* = 0.33, Amp: 0.90; adaptation: *M* = 0.11, *SD* = 0.46, Amp 0.78; difference: *F*_3,174_ = 9.48, *p* < .001). Further evaluation showed that the central tendency shifted, so that clockwise-drifting stimuli were now more likely to be judged as stationary (*F*_1,174_ = 9.79, *p* = .002) after adapting to a stimulus rotating clockwise. We also observed a significant increase in the standard deviation, so that a broader range of stimuli were associated with perceptual stasis (*F*_1,174_ = 20.45, *p* < .001).

#### Co-ordinate mapping

We then examined whether the effect of motion adaptation was dependent on the location of the test stimulus. Results showed that adaptation had a clear impact on the central tendency of retinotopic tests (*M* = 0.26, *SD* = 0.45, Amp: 0.85; baseline vs retinotopic: *F*_1,168_ = 64.53, *p* < .001) and also a small and opposite impact on the non-specific tests (*M* = −0.04, *SD* = 0.36, Amp: 0.93; baseline vs non-specific: *F*_1,174_ = 5.81, *p* = .017). Comparing test locations showed a large dissociation in central tendency between the test locations (retinotopic vs nonspecific: *F*_1,172_ = 82.25, *p* < .001).

### Discussion

Results of Experiment 5 show a retinotopic motion aftereffect and a clear dissociation between the retinotopic and non-retinotopic test locations.

## General Discussion

Our results suggest that causality aftereffects are not retinotopically-specific. We first showed that the boundary between subjectively causal and non-causal impressions is different to the point of subjective causality (Experiment 1). This revealed ambiguity about how to measure and interpret the effect of adaptation on subjective causality judgements, possibly due to a range restriction in previous studies. Measuring a broader range of stimuli (Experiment 2) verified that causality adaptation studies have failed to test important stimuli for eliciting causal impressions. We verified the claim that adaptation to contact-launch stimuli can change the point of subjective causality (Rolfs et al., 2013), but importantly, this effect did not appear to be location-specific (Experiment 3). We then ruled out the possibility that this spatial independence was due to using a neutral stimulus (Experiment 4) and demonstrated the validity of our general approach by measuring the retinotopic motion adaptation aftereffect (Experiment 5). In concert, these results directly repudiate an exclusively retinotopic effect of causality adaptation and suggest that the brain is not hardwired to encode visual causality through low-level, retinotopically mapped neural populations.

Causal impressions can be described for launch events across a range of spatial correspondences between two stimuli. In Experiment 1 we measured causal impressions for stimuli with spatial coincidence or overlap, and the resulting response distribution can be described by a cumulative Gaussian function. The central tendency of this function describes the degree of spatial coincidence at launch for which an observer is equally likely to report a causal launch or a non-causal pass. We also showed that these same data could be well described by a raised Gaussian distribution, with the central tendency of this function corresponding to the stimulus most likely to produce a causal impression. The ambiguity about which function most appropriately describes causal impressions resulted from a limited range of spatial correspondence (launch stimuli without spatial coincidence were not measured) between stimuli during a launch event.

We further showed that previous decisions impacted subsequent decisions. When we analysed data according to whether the previous response was a causal launch or a non-causal pass, we observed a change in central tendency of the cumulative Gaussian function, but not in the raised Gaussian function, even though the two functions were fit to the same data. This observation, that prior decisions can influence central tendency estimates depending on which model is fit, has important implications for measuring and interpreting adaptation aftereffects across a limited range of launch stimuli. Whether this effect of previous decisions represents a simple response bias, altered expectations, or a change in the perception of subsequent test events is unclear.

In Experiment 2 we collected baseline data across a broader range of launch stimuli, which verified that causal impressions were best described by a raised Gaussian distribution rather than a cumulative Gaussian distribution. Our results at baseline also showed that causal impressions can arise from stimuli with no spatial coincidence (described as ‘action at a distance’, Michotte, 1963, p.p. 99-100). Causal impressions were tightly constrained by the degree of spatial correspondence at the time of launch — too much overlap and too much separation each resulted in a reduction of the impression of causal launching. This result suggests that changes in the point of subjective causality (PSC) is the more appropriate measure of an adaptation aftereffect compared to the point of subjective equality (PSE). We then showed that adaptation to a contact-launch stimulus changes the PSC of the Gaussian distribution, confirming the presence of a causality aftereffect (Rolfs et al., 2013).

Experiment 3 showed that causal impressions were overall reduced after adaptation, consistent with findings from previous research (Rolfs et al., 2013; Burr; but see Arnold et al., 2015), but a significant adaptation aftereffect was observed in both retinotopic and non-retinotopic co-ordinates, and the aftereffect was not significantly different between these test locations. We suspected that the adapting stimulus might be too similar in its spatial correspondence to the stimulus most likely to produce a causal impression (i.e., a ‘neutral’ stimulus), and this might explain why the adaptation protocol reduced subsequent judgements of causality or failed to identify a co-ordinate mapping effect.

In Experiment 4 we adapted participants to action at a distance: an event in which one object appears to launch a second object, but the two never have spatial coincidence. This adapting stimulus also produced a reduction in reported causality for overlapping stimuli, but importantly, test events with spatial separation became *more likely* to result in causal impressions, and this effect was again location independent. Together, these findings strongly contradict the idea that launch adaptation reduces causality perception through low-level, retinotopically-mapped causality-detection channels in early visual cortex. Indeed, we found an increase in judgements of causality for stimuli with spatial disunity in both retinotopic and non-retinotopic co-ordinates.

In Experiment 5, we further demonstrated the validity of our approach by measuring a retinotopically-mapped motion aftereffect, thus ruling out the possibility that our Experimental design was unable to detect location-dependent effects of causality adaptation.

The present study provides counter-evidence for the claim that causality adaptation is exclusive to retinotopically-mapped regions of visual space. We argue that conclusions of spatially localised causality aftereffects rely, in part, on the misapplication of important aspects of psychophysical modelling (see Experiment 1). When accounting for the full range of the spatial constraints of causal impressions (Experiment 2), and then measuring spatial localisation effects (in Experiments 3 & 4), we found that causality aftereffects were location independent, and that our Experimental design was appropriate for detecting such effects (Experiment 5). We suggest that causality aftereffects do not reflect changes in low-level visual processes, like those observed in the aftereffects of motion and colour and orientation.

Other aspects of our study further support our conclusions. Our study tested a total of 70 naïve observers, 30 of whom were tested specifically for spatially localised effects of causality adaptation. Contrast our sample with the four participants in Rolfs and colleagues (2013) critical Experiment 2. Consider also that in their Experiment 1 they found support for a non-retinotopic aftereffect with a sample size of eight. This leaves open the possibility of a type 2 error in their conclusion that causality aftereffects arise from the adaptation of retinotopically mapped causality detectors. Additionally, Rolfs and colleagues’ data are (to our knowledge) the only data reported, to date, demonstrating retinotopic specificity of causality adaptation. The present study, on the other hand, is consistent with the results of their first Experiment, and also a previous study by some of the present authors (see Arnold et al., 2015), making this study the third demonstration that causality aftereffects can be location independent. Rolfs and colleagues did, however, test for adaptation using 320 adapting events compared to our 18 initial and 6 top-up adapting events. Perhaps this gave them greater power to detect retinotopic effects. Equally possible is that other (more basic) aspects involved in causal impressions became adapted at the critical location (e.g., contrast sensitivity, or motion adaptation) or produced an expectation effect between test locations. Regardless, our data strongly suggest that causality aftereffects occur in spatially global co-ordinates.

## Conclusion

While our results cannot directly rule out that causality is itself a perceptual representation (rather than a strong and automatic inference), our data appear to indicate that expectations play an important role in response changes in a causality aftereffect study.

